# “Genome-wide population diversity in *Hymenoscyphus fraxineus* points to an eastern Russian origin of European Ash dieback”

**DOI:** 10.1101/154492

**Authors:** Jørn Henrik Sønstebø, Adam Vivian-Smith, Kalev Adamson, Rein Drenkhan, Halvor Solheim, Ari M. Hietala

## Abstract

European forests are experiencing extensive invasion from the Ash pathogen *Hymenoscyphus fraxineus*, an ecological niche competitor to the non-pathogenic native congener *H. albidus*. We report the genome-wide diversity and population structure in Asia (native) and Europe (the introduced range). We show *H. fraxineus* underwent a dramatic bottleneck upon introduction to Europe around 30-40 generations ago, leaving a genomic signature, characterized by long segments of fixation, interspersed with “diversity islands” that are identical throughout Europe. This means no effective secondary contact with other populations has occurred. Genome-wide variation is consistently high within sampled locations in Japan and the Russian Far East, and lack of differentiation amongst Russian locations suggests extensive gene flow, similar to Europe. A local ancestry analysis supports Russia as a more likely source population than Japan. Negligible latency, rapid host-range expansion and viability of small founding populations specify strong biosecurity forewarnings against new introductions from outside Europe.

## Introduction

Establishment likelihood and impact of an invasive species can be often associated by the ecological similarity of species assemblages at different locations (Paini et al. 2016), the presence of specific ecological drivers (Dawson et al. 2017), the number of introductions and the viability of small populations (Blackburn et al. 2011, Fauvergue et al. 2012). Understanding evolutionary processes associated with biological invasions such as loss of genetic variation due to reduction in population size at the founder event and change in selection pressure in the new environment is challenging (Zhan et al. 2012), but necessary to comprehend the historical process and future trajectory of the invasion. Population genomic analyses use genome scale sequence data to enable the study of these evolutionary processes (Grunwald et al. 2016). Comparing the genome wide genetic structure and diversity between native and introduced populations enables detailed studies of the genetic and evolutionary aspects of the invasion. Principal questions to address are the number of introduction events, the mode of establishment, the initial effective population size, the effects of the founder event on the genetic structure and diversity, and the differences in adaptive potential between the native and the introduced populations.

Dieback of European Ash (*Fraxinus excelsior* L.) is one of the most important fungal diseases on trees in Europe as it threatens a keystone tree species and associated forest biodiversity. The causative ascomycete *H. fraxineus* appears to have been introduced to Europe from Asia, where similar parallel species assemblages exist. In Asia *H. fraxineus* is reported as extensive leaf colonizer and notably a weak pathogen of *Fraxinus mandshurica* and *F. rhynchophylla* (Zhao et al. 2013, Zheng and Zhuang 2014, Drenkhan et al. 2017). In contrast to its role as a leaf colonizer and weak pathogen on Asian ash species, *H. fraxineus* is able to grow into shoots and branches of European ash and eventually kill the tree. Young trees die within a few years of infection while older trees often become chronically infected (Kowalski and Holdenrieder 2009, Skovsgaard et al. 2010). European Ash has also been historically associated with a non-pathogenic endophyte *H. albidus* (Baral and Bemmann 2014, Baral et al. 2014), a species experiencing invasion-induced range contraction through direct niche competition (McKinney et al. 2012).

The *H. fraxineus* invasion relies on a high sexual ascospore production (Cross et al. 2017), consistent with propagule pressure theory (Lockwood et al. 2005). The life cycle can be completed annually via the formation of apothecia on leaf petioles and rachis debris left on the forest floor (Kowalski and Holdenrieder 2009, Timmermann et al. 2011). Synchronous ascospore discharge (Hietala et al. 2013) appears to coordinate host challenge and facilitate long distance dispersal, as proposed for other ascomycetes (Roper et al. 2010). While *H. fraxineus* is an outcrossing fungus with equal representation of the two mating types in Europe (Gross et al. 2012), it additionally produces asexual spores (Kowalski and Bartnik, 2010; Fones et al. 2016), but as no clonal populations occur these have been proposed to function primarily as spermatia (Gross et al. 2012). Gene introgression from native species may facilitate adaptation of invasive plant pathogens (Gonthier and Garbelotto 2011), yet the population structures and relationships between *H. fraxineus* and the native competitor *H. albidus* have not been characterized at genome-wide scales.

Ash dieback symptoms were first recorded in Poland in the early 1990s (Przybył 2002), and the first European record of *H. fraxineus* is from Estonia from 1997 (Drenkhan et al. 2016). Since 2001, extensive spread has been observed throughout Europe and into Scandinavia (McKinney et al. 2014). The disease front advanced in 1992-2008 from eastern Poland to Switzerland with an annual dispersal rate of 75 km per year (Gross et al. 2014). Genetic diversity and population structure studies have provided some broad indications about the spread of *H. fraxineus* in Europe (Rytkönen et al. 2011, Bengtsson et al. 2012, Gross et al. 2012, Kraj et al. 2012, Gross et al. 2014, Burokiene et al. 2015). Schoebel et al. (2017) also recently characterized the mitovirus population associated with *H. fraxineus* in Europe, showing the presence of two ancestral types, perhaps indicative of two founding parents. Together these prior studies provide a narrative that for a lack of population structure and low European allelic richness: as compared to Japanese populations on *F. mandschurica*, the European population was established by two individual strains in eastern Europe (Gross et al. 2014), although few informative microsatellite loci have been studied.

Here we report, alongside McMullan et al. (submitted), who sequenced and assembled the *H. fraxineus* genome, the genome-wide diversity of multiple *H. fraxineus* populations in Europe, with reference sampling locations in Far Eastern Russia and Japan (Figure 1A,B), using densely spaced single nucleotide polymorphisms (SNPs). We support the conclusion of McMullan et al. (submitted) that only two haploid *H. fraxineus* individuals founded the European population, and show that this led to a genome-wide reduction in genetic diversity compared to the native populations. We show that the bottleneck experienced in the European population drastically reduced genetic diversity and caused fixation of large swathes of the genome, which are detected in all European samples. Importantly we use this data, along with the allelic richness and genetic differentiation in the Russian Far East and Japanese populations, to provide information about the potential for local adaptation, and estimate the time since the introduction to Europe, and conclude, through a local ancestry analysis, that the Russian Far East is a likely source for the invasive population.

**Figure 1:**
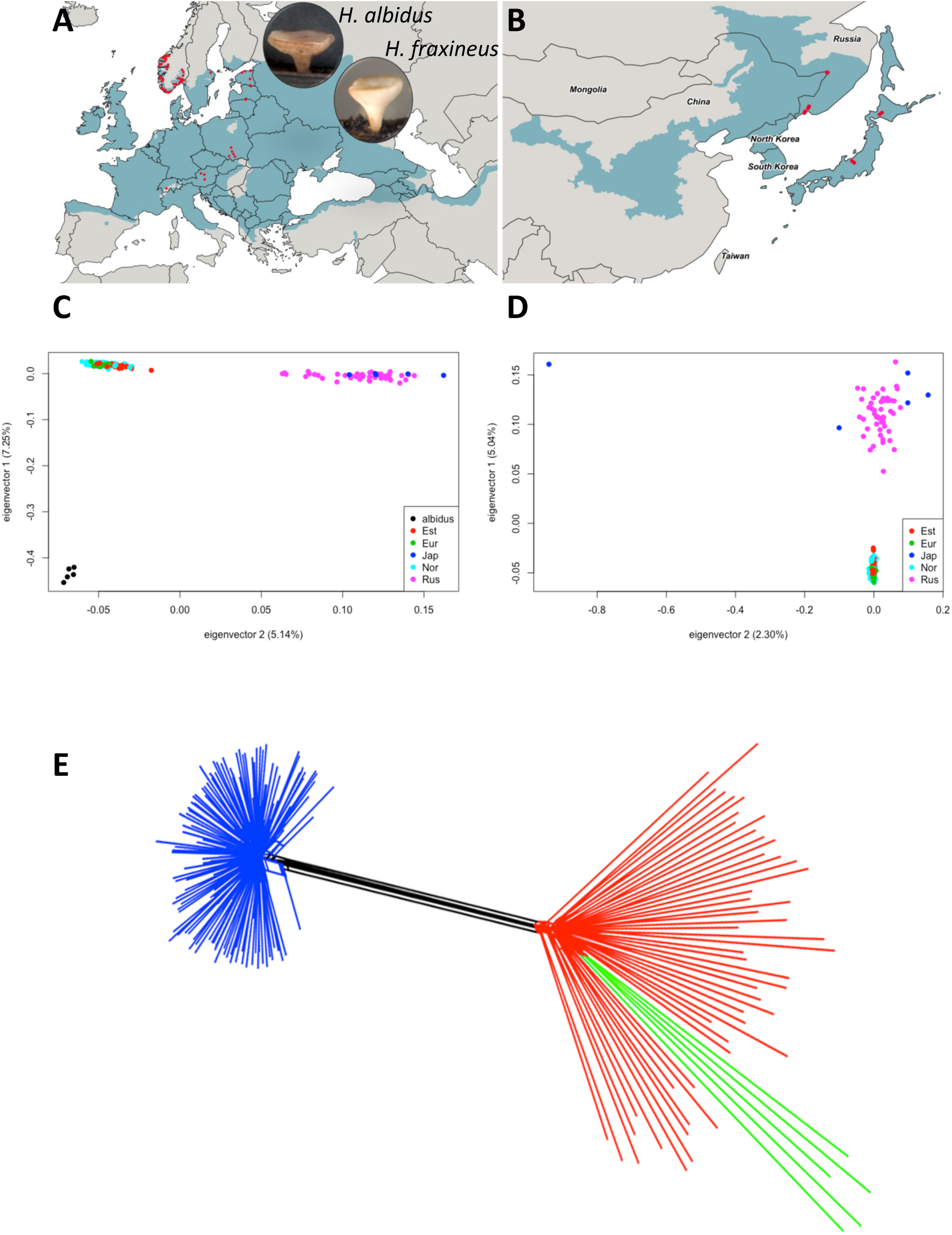
Sampling locations and relationship among isolates of *Hymenoscyphus fraxineu*s and *H. albidus*. Map of sampling locations in Europe (A) and Asia (B). Principle component analysis of *H. fraxineus* and *H. albidus* (C) and *H. fraxineus* only (D). The percentage of variance held by each eigenvector is written on the axis. The first eigenvector splits *H. albidus* and *H. fraxineus*, while the second splits the Asian and the European populations of *H. fraxineus*. The SplitsTree, neighbor-net analysis of *H. fraxineus* (E) shows the same pattern of relationship between the different populations as the PCA where the European samples (blue) are separated from the Russian (red) and the Japanese (green).

## Material and methods

### *Hymenoscyphus fraxineus* samples

Samples of *Hymenoscyphus fraxineus* were collected from Estonia (33 individuals), Norway (90) and the Russian Far East (51). Additional samples were obtained from Japan (5) and from other locations in Europe (14), including the holotype strain of *H. fraxineus* (see Supplementary Table 1). We analysed in total 193 *H. fraxineus* individuals. Estonia and Norway were specifically chosen as representative locations because of distinct disease histories. The first European record of *H. fraxineus* is from 1997 from Estonia (Drenkhan et al. 2016). In Norway, Ash dieback was first documented in southeastern parts of the country in 2008, after which the disease spread along the west coast at an annual rate of 51 km (Solheim and Hietala (2017); Figure 1A).

Sampling in Norway and Estonia was performed over several years, with Norway from 2008 to 2014 covering the disease fronts along the southern and western coasts (Supplementary Table1). Estonian sampling was conducted from 2010 to 2014, while the samples from the Russian Far East were collected in 2014 in three locations (Figure 1, Supplementary Table 1). The majority of Norwegian samples represent haploid monokaryotic isolates, with some obtained as a single spore cultures (SSC) from ascocarps, and some as a single hyphal isolates from mycelial *in vitro* cultures derived from necrotic shoot lesions (Supplementary Table 1). All Estonian and Russian samples were ascocarps, meaning they are heterokaryotic and contain the parental strains and sexual progeny. In addition to the *H. fraxineus* samples we also included five *H. albidus* isolates from Norway and Switzerland (Supplementary Table 1).

### ddRAD analysis

DNA samples from all Estonian and the Russian Far East ascocarps were extracted using the E.Z.N.A Fungal DNA Mini Kit (Omega Bio-Tek Inc., Norcross, GA, USA). Norwegian DNA samples were isolated using the Easy-DNA protocol No. 8 for Mouse Tails (Thermo Fisher Scientific). DNA quantity was assessed with the Nanodrop 2000 (Thermo Fisher Scientific), and/or the Qubit DNA Broad Range Sensitivity assay (Thermo Fisher Scientific). Of the five Japanese isolates, DNAs from three Japanese isolates were directly amplified from mycelia using the Repli-G Single Cell kit using whole genome amplification (P/N 150343; Qiagen; Supplementary methods and Supplementary Table 1).

All samples were pretested by species-specific ITS PCR and Sanger sequencing using HFrax-F and HFrax-R primers as described previously (Drenkhan et al. 2016, Cross et al. 2017). All the samples were positive and provided typical alignments to *H. fraxineus* except for two samples from Russia and one Estonian sample (IDs 4054; 4142 and 4137 respectively), which were excluded later after ddRAD sequencing, since these samples were contaminants from unrelated species (Supplementary Table 1).

Each DNA sample consisted of a normalized input of 200 ng, which was prepared in 30 μl of nuclease free water. A single digestion and ligation reaction comprised the DNA with *PstI-HF* and *MspI* (New England Biolabs; R3140L and R0106L respectively) and the ddRAD ligation adapters compatible to the restriction sites together with 20 units of each restriction enzyme, 100 units of T4 DNA ligase and 1 μl of 10 mM ATP, in a total volume of 40 μl. Reactions were pooled and purified in the subsequent steps with 1.1x volumes of AmpureXP (Beckman Coulter, Inc.). Samples with compatible barcodes were size fractioned together to 183 to 383 bp with the Pippin Prep and a 2% agarose gel cassette (2010CDS) using marker B (Sage Science, Beverly, MA, USA). Size selected libraries were further purified with AmpureXP and then amplified in a total volume of 260 μl with Platinum PCR Super Mix High Fidelity polymerase (Thermo Fisher), 10 μl each of the P and A1 primer (20 μM) as described in Ion Xpress library amplification (Thermo Fisher Scientific; Vivian-Smith and Sønstebø, 2017). Libraries were qualified and quantified using a Bioanalyzer 2100 High Sensitivity DNA chip (Agilent) before appropriately diluting the sample and populating Ion Sphere Particles (ISPs) with template. Emulsion PCR and enrichment of ISPs were either performed using the Ion One Touch 2 system using OT2 reagents (Thermo Fisher Scientific Publication Part Number MAN0007221 Rev. 2.0), or the Ion Chef using IC 200 reagents (MAN0007661 Rev. A.0). Sequencing was performed on an Ion torrent PGM sequencer using 316 chips and monitored with FastQC (Leggett et al. 2013).

### Bioinformatics

The ddRAD sequences were de-multiplexed using Torrent Server software suite 4.6. The sequences from the individual samples were then quality trimmed in CLC Genomic Workbench 8 (<u>www.qiagenbioinformatics.com</u>). Sequences passing this filter were mapped to the reference genome KW1 *Hf*v2 (Saunders et al. 2014, McMullan et al. submitted), using Novoalign (Novocraft Technologies, Selangor, Malaysia).

Multi-sample variant calling was performed with FREEBAYES (Garrison and Marth 2012), with --min-alternate-count 4 --min-alternate-fraction 0.2. Given the generally higher rate of errors in indels compared to SNPs in Ion Torrent data, we filtered out all the indels and kept only the SNPs in the further analyses. This was done using VCFFILTER in VCFLIB (<u>https://github.com/vcflib/vcflib),</u> where also only variants with a QUAL score above 30 were retained. SNPs present in less than 20% of the individuals were removed as well as SNPs with a minor allele frequency (MAF) of less than 1%. The MAF was set low since many of the loci that are monomorphic in Europe show variation in the Asian population. Increasing the limit of the MAF to 0.05 led to many of the Asian specific alleles being lost. Filtering of SNPs was performed in VCFTOOLS (Danecek et al. 2011). The SplitsTree (Huson and Bryant 2006) and the STRUCTURE v2.3.4 (Pritchard et al. 2000, Falush et al. 2003) analysis (see below) was performed on a thinned data set, at one marker per contig, using the thinning procedure in VCFTOOLS.

### Population genomic analysis

McMullan et al. (submitted) detected a highly reduced genetic diversity in European *H. fraxineus* throughout the genome compared with the Japanese population, suggesting that the European population was founded by a few individuals. In order to confirm this, we calculated the nucleotide diversity in the R-package PopGenome (Pfeifer et al. 2014) for each population, except Japan, for which too few isolates were available. The calculations were done in windows of seven SNPs with a step of five SNPs. The pattern of nucleotide diversity was also assessed with the calculation of nucleotide divergence π c in VCFTOOLS (Danecek et al. 2011). Here, π c was calculated over sliding windows of 20 kb with 5 kb steps. We combined all samples from Europe to contain the overall diversity in Europe and compared that with all Russian samples.

Tajima’s D was used to detect departures from the standard neutral model and calculated using VCFTOOLS (Danecek et al. 2011).

To further assess signature of selection, we used the Bayesian likelihood method implemented via reversible jump Markov Chain Monte Carlo in BayeScan (Foll and Gaggiotti 2008). In short, BayeScan estimates the probability that a locus is under selection by calculating a Bayes factor, which is simply the ratio of the posterior probabilities of two models (selection/neutral) in the given data. Bayes factors above 100 (log10 > 2) correspond to posterior probabilities between 0.99 and 1. BayeScan has been shown to detect most loci under strong selection, with low probability of detecting false positive (Narum and Hess 2011). The BayeScan analysis was performed separately in Europe and Asia. In Europe we divided the Norwegian population into an Eastern (early introduction) and a Western (later introduction) population and used these along with the Estonian population to detect selection (the groups are described in Supplementary Table 1). In Asia we used the three sampling locations as populations and used these to infer loci under selection.

The relationship among samples of *H. fraxineus* and *H. albidus* were investigated with principle component analyses (PCA) in the R package SNPRelate (Zheng et al. 2012), using only loci with a level of linkage disequilibrium r smaller than 0.2. We also performed a phylogenetic network analysis using the Neighbor-Net algorithm in SplitsTree 4.0 (Huson and Bryant 2006). Concatenated SNPs were extracted from the VCF file using PGDSpider (Lischer and Excoffier 2012). SplitsTree was performed using uncorrected P distances and the heterozygous sites were treated as average between the two alternative states. In both the PCA and SplitsTree analysis we included *H. albidus* as an outgroup, but also to examine signs of hybridization between the two species. The genetic differentiation between the two species and among sampling locations within *H. fraxineus* was also investigated with sliding window FST analyses along the genome (100kb windows, 50kb steps) using VCFTOOLS.

The genetic structure of *H. fraxineus* was assessed using STRUCTURE v2.3.4 (Pritchard et al. 2000, Falush et al. 2003). In STRUCTURE, individuals are placed into K clusters that are in H-W and linkage equilibrium, and have distinctive allele frequencies without imposing *a priori* population information. K was chosen in advance, but was varied between 1 and 6. For each K value the analysis was iterated 20 times using an initial burn-in of 50 000 cycles, and then another additional 50 000 cycles. Individuals may have membership in several clusters (indicating admixture) with the membership coefficient summing to 1. STRUCTURE output, Pr(X|K), can be used as an indication of the most likely number of genetic groups and the level of admixture between groups. However, as indicated in the STRUCTURE documentation and according to Evanno et al. (2005), Pr(X|K) plateaus or slightly increases after the best K is reached. Thus, following Evanno et al. (2005), deltaK was calculated and used to evaluate the most probable genetic structure. Delta K was calculated using STRUCTURE HARVESTER (Earl and vonHoldt 2012) and runs with identical K were combined with CLUMPP (Jakobsson and Rosenberg 2007). Finally PCAdmix was used to infer local ancestry (Brisbin et al. 2012). This approach relied on phased haplotypes derived from both reference panels and the admixed individuals. We used Beagle 4.0 (Browning and Browning 2007) to phase the data from ascocarps, without imputation. This was done individually for each population. The local ancestry was inferred with PCAdmix using windows of 10 SNPs. For each window, the distribution of individual scores within a population is modeled by fitting a multivariate normal distribution. These distributions are then used to compute the likelihood of each score belonging to either Japan or Russia. Local ancestry assignments were determined using a 0.9 posterior probability threshold for each window. Samples having probability of belonging to a reference of < 0.9 were recorded as missing ancestry.

## Results

ddRAD sequencing provided an almost uniform coverage across the *H. fraxineus* genome (Supplementary Figure 1), and an average of 0.86% total of the genome was consistently resequenced at each ddRAD loci for each sample. Small tracts of the genome that represented specific sequence compositions that did not cut with *PstI* and *MspI* were under-sampled, the largest being around 250 kbp (Supplementary Figure 1; scaffold 13). The amounts of reads that mapped to the genome also varied between samples (Supplementary Figure 2), but in general, 90% of the reads in each of the European and Japanese mycelial cultures mapped to the Hfv2 genome (McMullan et al., submitted). Lower levels of the *H. fraxineus* ascocarp reads mapped and we observed a larger variation in the percentage of mapped reads per individual. For the *H. albidus* cultures around 75% of the reads mapped to the *Hfv2* reference genome (Supplementary Figure 2). BLAST searches from the unmapped reads from the ascocarps detected contaminants such as *Pseudomonas aeruginosa*. The variant calling with FREEBAYES detected in total 71335 variable sites of which 49019 were SNPs. After removing SNPs with a maximum amount of missing data of 0.2 (44564 left) and a minimum allele frequency < 0.01, 15181 SNPs were left for further analyses.

### *H. fraxineus* and *H. albidus* are well separated species

Principle component analysis separated *H. fraxineus* and *H. albidus* along the first eigenvector, which held 7.25 percent of the variance (Figure 1C). The clear separation between *H. fraxineus* and *H. albidus* was also evident in the SplitsTree analyses where all the *H. fraxineus* individuals were separated from *H. albidus* with a single long branch containing both the Swiss and Norwegian isolates (Supplementary Figure 3). Similarly the FST analysis showed a high differentiation between European or Asian *H. fraxineus* and *H. albidus*: mean pairwise F_ST_ for *H. albidus* and European or Asian *H. fraxineus* were 0.767 (sd 0.208) and 0.420 (sd 0.243), respectively. This pattern was consistent across all scaffolds (Supplementary Figure 4) suggesting that introgression between the two species has not taken place or the level is too low to be detected by these analyses. However, European *H. albidus* is additionally differentiated from *H. fraxineus* based on a preliminary genomic resequencing analysis and mapping of *H. albidus* sequencing reads to *Hfv2* scaffolds (Vivian-Smith, pers. comm.). Comparisons of the total GC content and a K-mer analysis, show consistent differences between the two species (Supplementary Figure 5). Therefore, the population structure analyses, together with preliminary genomic data, unambiguously resolves *H. albidus* into a separate species.

### Population structure analyses show clear division of native and introduced *H. fraxineus* populations, with a variable level of structuring between two biogeographic regions

The genome-wide PCA divided the Asian and the European samples of *H. fraxineus* into two clearly separated groups along the first eigenvector, while the second eigenvector mainly showed variation in the Asian individuals (Figure 1D). In comparison, the single-scaffold PCA also separated the European and the Asian samples along the first axis for all scaffolds (Supplementary Figure 6), but their differentiation was much weaker than in the genome-wide PCA (Figure 1D). The clear distinction between the Asian and the European populations was also found in the neighbor-network SplitsTree analysis, where the Asian and European groups connected with a few long parallel branches (Figure 1E). Strains from Japan were embedded in the Russian group, but had longer branches than the samples from Russia, which may be due to origin at widely different locations in Japan. In the European group, all isolates were largely connected at one node potentially representing a single common ancestor at the foundation of the European population, a finding which is reinforced when the Neighbor-Net split decomposition analysis is performed with only the European isolates (Supplementary Figure 7). STRUCTURE analysis with K = 2 also realized that two genetic groups were almost exclusively divided between Asia and Europe (not shown).

Separation between Japanese and Russian populations was detected in the PCA analysis when STRUCTURE was performed on only the Asian samples (Figure 2). With K=2 the Asian samples divided into a Japanese and a Russian group with very little admixture. With K=3 a third group was observed that had a slightly variable frequency across the three Russian survey locations. Within Europe, STRUCTURE detected two groups that were not related to geography or the time since local introduction. These groups are suspected to represent the two historical parental haplotypes that founded the European population, with each group displaying the residual allele frequency of the two original parental haplotypes.

**Figure 2:**
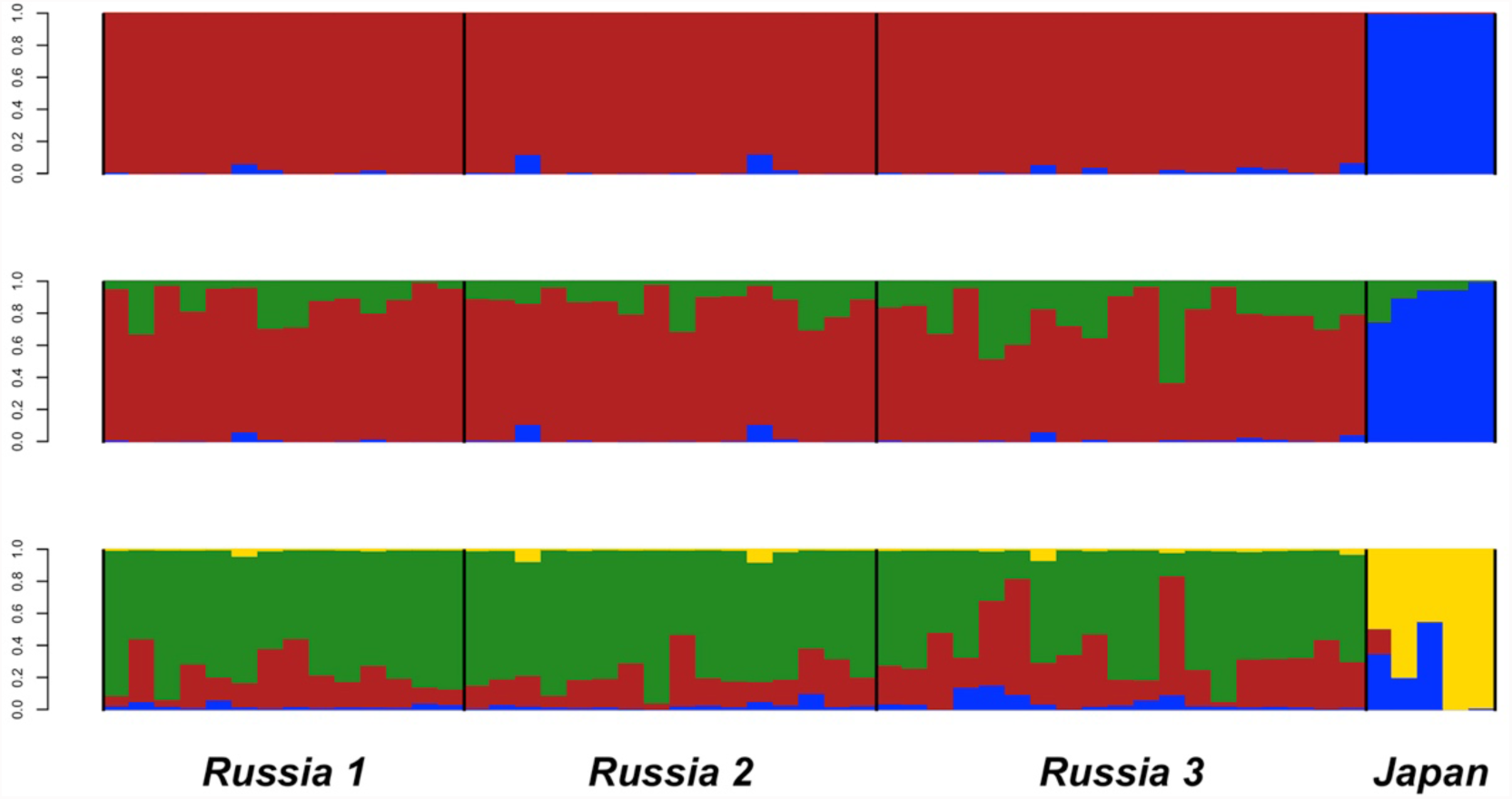
Population structure of Asian *Hymenoscyphus fraxineus*. Structure analysis of the Asian *H. fraxineus* individuals shows the membership in different genetic groups at K=2 (top), K=3 (middle) and K=4 (bottom).

### European populations show a fixation signature across the majority of scaffolds

The magnitude of the genome-wide reduction in nucleotide diversity varied considerably amongst scaffolds, but an identical pattern was discovered throughout all three European populations (Figure 3). In contrast, populations from Russia had a high genome-wide nucleotide diversity (Figure 3). We considered that the regions of low nucleotide diversity were the tracts of the genome where fixation event(s) had occurred and termed the regions with higher nucleotide “diversity islands”. Several scaffolds appeared to have entirely preserved their former diversity and escaped fixation (scaffolds 13 and 15; Figure 3). Other scaffolds have almost been exclusively fixed (scaffolds 14, 16 and 20). Similar pattern was observed in the π c calculated in windows of 20 kb (Supplementary Figure 8). However, we did observe several small diversity islands within these chromatids. These were either representative of a close and coincident double recombinant event, or a set of sequential inter-generational recombinant events that then led to fixation on either side of the diversity island. The proportion of smaller diversity islands throughout the genome (Figure 3), along with the near complete fixation of entire chromatids, further denote that the fixation event occurred in a succinct and sequential manner after immigration. Importantly, this occurred prior to the European *H. fraxineus* isolate becoming numerically superior and spreading throughout the whole European region, thus leaving a fixation signature homogenous to the whole of Europe.

**Figure 3.**
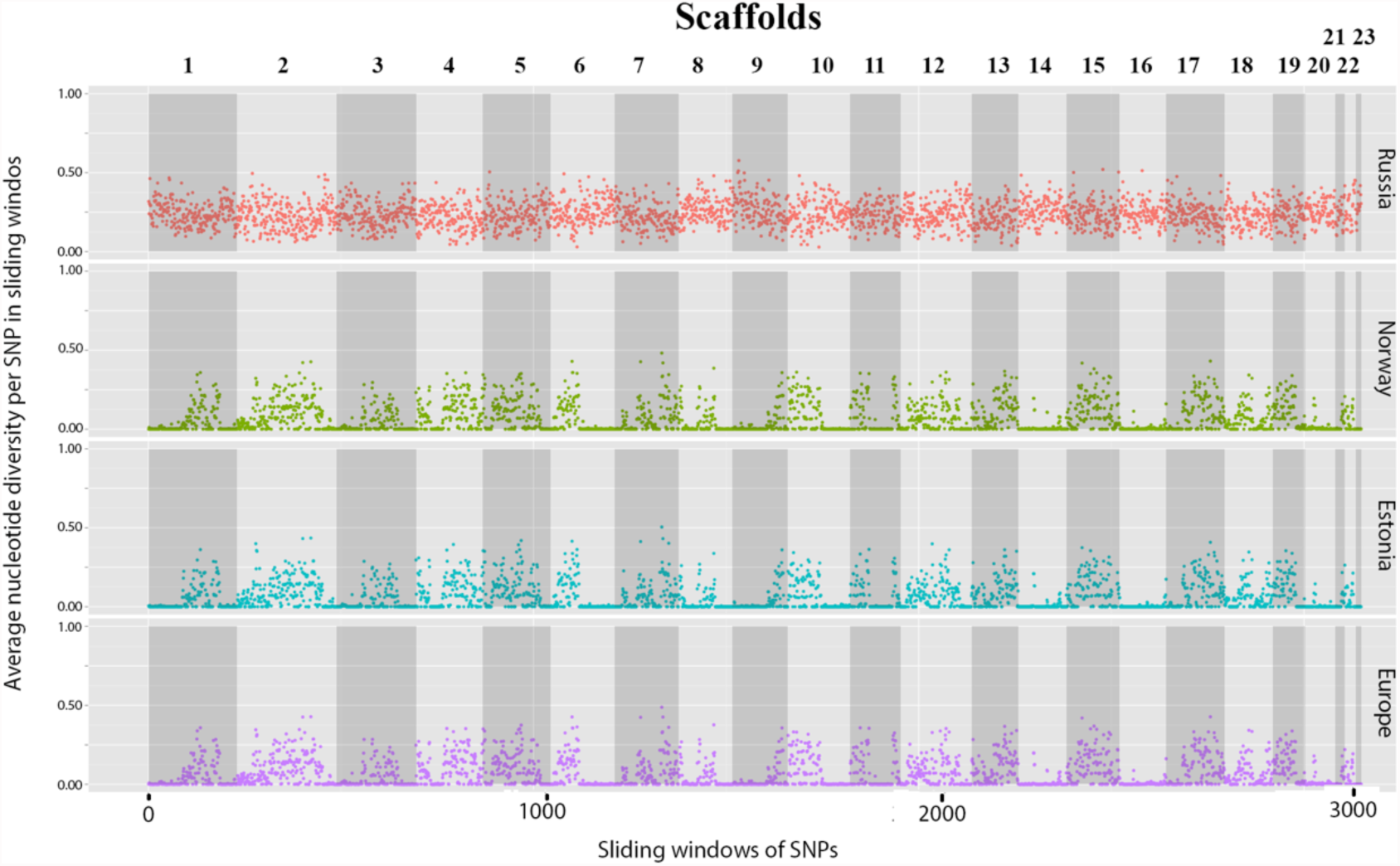
Genome-wide diversity of *Hymenoscyphus fraxineus*. Nucleotide diversity per SNP calculated in sliding windows of seven SNPs with five SNPs jumps in the Russian Far East, Norwegian, Estonian and remaining European individuals.

The pattern of segmentation for the fixed tracts and diversity islands, which is expected to be a signature of a bottleneck from the founding population, yields information on the establishment phase and breeding behavior. Therefore, each of the fixed segments, and diversity islands was assessed for the length of that tract and their occurrence frequencies at by calculation in R (Supplementary Methods). The average length of the fixed blocks was 68 Kbp, with a maximum spanning length of 1.13 Mbp. Likewise, the average length of the polymorphic blocks was 54.6 Kbp, with a maximum spanning length of 245 Kbp (Supplementary Figure 9). It follows that the timing since the fixation event can be approximated based on the level of recombination and the subsequent fragmentation of the genome into non-variable segments (Stukenbrock et al. 2011, Stukenbrock et al. 2012), even though these segments will have aggregated to a certain extent. Estimates of recombination rates for ascomycetous fungi range from 46 cM/Mb in an ancestral type of *Zymoseptoria* (Dothideomycetes), to between 86 and 128 cM/Mb for *Botrytis cinerea* (Helotiales; (Amselem et al. 2011, Van Kan et al. 2017)). Taking the mean 68 Kbp size of the non-variable segments of the European *H. fraxinus* as a chronicle of the historical recombination process since the fixation event, it can be postulated that if a recombination event occurred every 2.2 Mb per generation (Stukenbrock et al. 2012), then the time since fixation is 32 generations/years. Taking the mean of 98.6 cM/Mb for *Botrytis cinerea* as a proxy leads to an estimate of around 14 years. Likewise, estimates from the variable segments provide estimates of 40.3 years, and 18.6 years given the recombination rates of 46cM/Mb and 98.6 cM/Mb as observed in *Zymoseptoria* sp. and *Botrytis cinerea* respectively. While this data does not identify the contribution of ancestral haplotypes to the non-variable segments, the data provided an opportunity to estimate the total proportion of the genome under fixation. This stands at 55.2%, and indicates some particular limitations on inter-generational breeding. Importantly, the parental strains appear to have been labile and characteristic of an annual life cycle, and the two founding individuals did not provide a continued contribution to the next generation, so a portion of their genetic contribution was lost through the fixation event.

### European genetic diversity is bottlenecked but the potential exists for local adaptation

McMullan et al (submitted) suggested, based on the diversity in effector genes that adaptive potential was still present in Europe. In order to compare the level of local adaptation in Europe and Asia we performed outlier analyses with BAYESCAN for the set of sampling sites in Europe (Estonia, eastern Norway and western Norway) and for the three Russian sampling sites. The populations in Europe and Russia had a similar genetic differentiation (Supplementary Figure 10), but the populations in Europe covered a larger biogeographical distance with different climatic zones. We detected 34 loci, dispersed across several scaffolds, with significant outlier behavior in Europe (corresponding to P value = 0.05). Of these 29 had P-values < 0.01, compared with only five in Russia, suggesting that local adaptation has already occurred since the introduction to Europe.

The average Tajima’s D for each scaffold varied considerably in Europe, from −1.35 within scaffold 14 to 0.91 in scaffold 15 (Figure 4). A positive Tajima’s D in single genetic regions is evidence for heterozygous sites having a selective advantage, while negative values suggest directional selection for specific allele. If, however, the majority of the genome has either a negative or a positive Tajima’s D value, the most probable explanation is that the population underwent a recent expansion or a bottleneck, respectively. In general, Tajima’s D was negative for the scaffolds with low nucleotide variation and for the large blocks fixed for one allele, while it was positive for the polymorphic scaffolds (Figure 4). The positive Tajima’s D at the polymorphic loci probably reflects the founder event and the loss of rare alleles upon this event. The negative value in the monomorphic scaffolds may be a result of new mutations with low frequency, reflecting the recent increase in population size.

**Figure 4.**
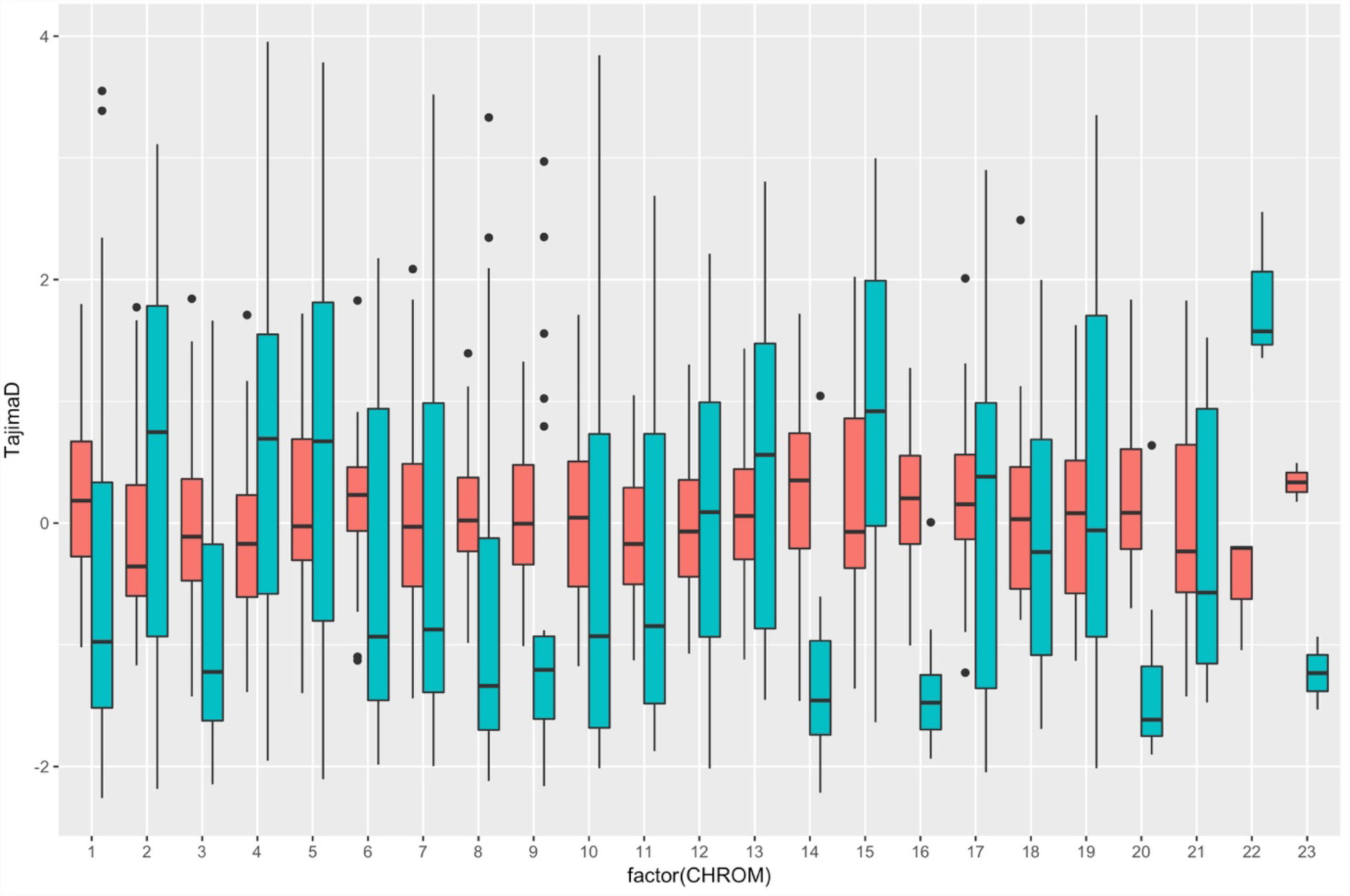
Tajima’s D for each scaffold. The Tajima’s D was calculated for the combined *H. fraxineus* from Asia (red) and Europe (green).

### European *H. fraxineus* comes from a locality near the Russian Far East

The population structure detected in Europe may have been amplified by the strong genetic drift and extensive linkage disequilibrium experienced by the European population of *H. fraxineus* during and after the founder event as is evident from the genome wide loss in genetic diversity (Figure 3). Also, although the European group appeared highly diverged from the Asian populations in both the PCA, SplitsTree and the STRUCTURE analysis, only a total of 784 alleles were private to Europe, while 1178 were private to the five Japanese samples and 8244 alleles were private to Asia (Figure 5). In total 978 alleles were shared among Norwegian, Estonian and Russian samples, but absent in Japan, while 77 alleles were found in only Norway, Estonia and Japan. This suggests a closer relationship between Europe and Russia, than Europe and Japan.

**Figure 5.**
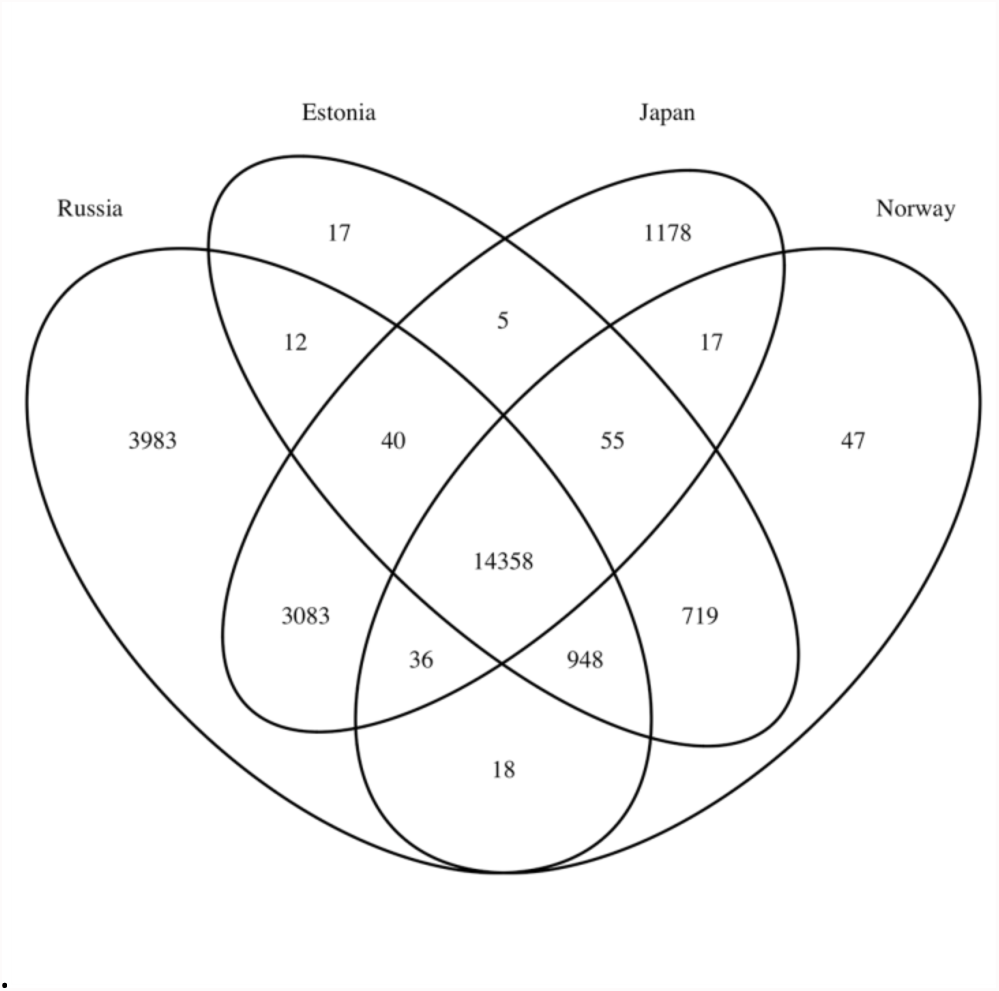
Distribution of alleles among sampling locations of *H. fraxineus*. The Venn diagram shows the distribution of alleles with frequency above 0.05 in *H. fraxineus* from Russia, Estonia, Japan and Norway.

In strong founder events, the low effective population size will affect the level of genetic drift and linkage disequilibrium resulting often in large changes in allele and haplotype frequencies. In order to infer the ancestry of the European samples we performed a local ancestry assignment using PCAdmix with Russia and Japan as ancestral groups. The local ancestry assignment was performed with PCAdmix in windows of ten SNPs along the genome for Norway and Estonia separately shows that a much closer Russian ancestry of the European samples is observed (for both Norwegian and Estonian), than towards the Japanese isolates (Figure 6). This corresponds well with the number of shared alleles found amongst these populations. Since the ancestral admixture assignment in PCAdmix examines the ancestry of small windows of SNPs it will be less affected by the changes in haplotype frequencies resulting from the genetic drift and linkage disequilibrium in the founding population. However, the results from the PCAdmix ancestry analysis were not homogenous throughout the genome. While the majority of windows had highest probability of a Russian origin, around 10% had Japan as the likely origin. These included portions of scaffolds 17, 9, 8, 6, 4, 3 and 2. Since the number of individuals used in the analysis was much higher for Russia than Japan we performed an analysis with similar numbers of random Russian haplotypes as the Japanese (5). This increased the Japanese ancestry in the genome to around 15%. Even though the fixed tracts of the European genome have not yet been assigned a parental haplotype group, smaller amounts of Japanese origin are likely explained by an ancestral introgression event in the native population preceding the introduction to Europe, or alternatively that the European ancestral population is located slightly closer to Japan than those sites sampled in Russia.

**Figure 6:**
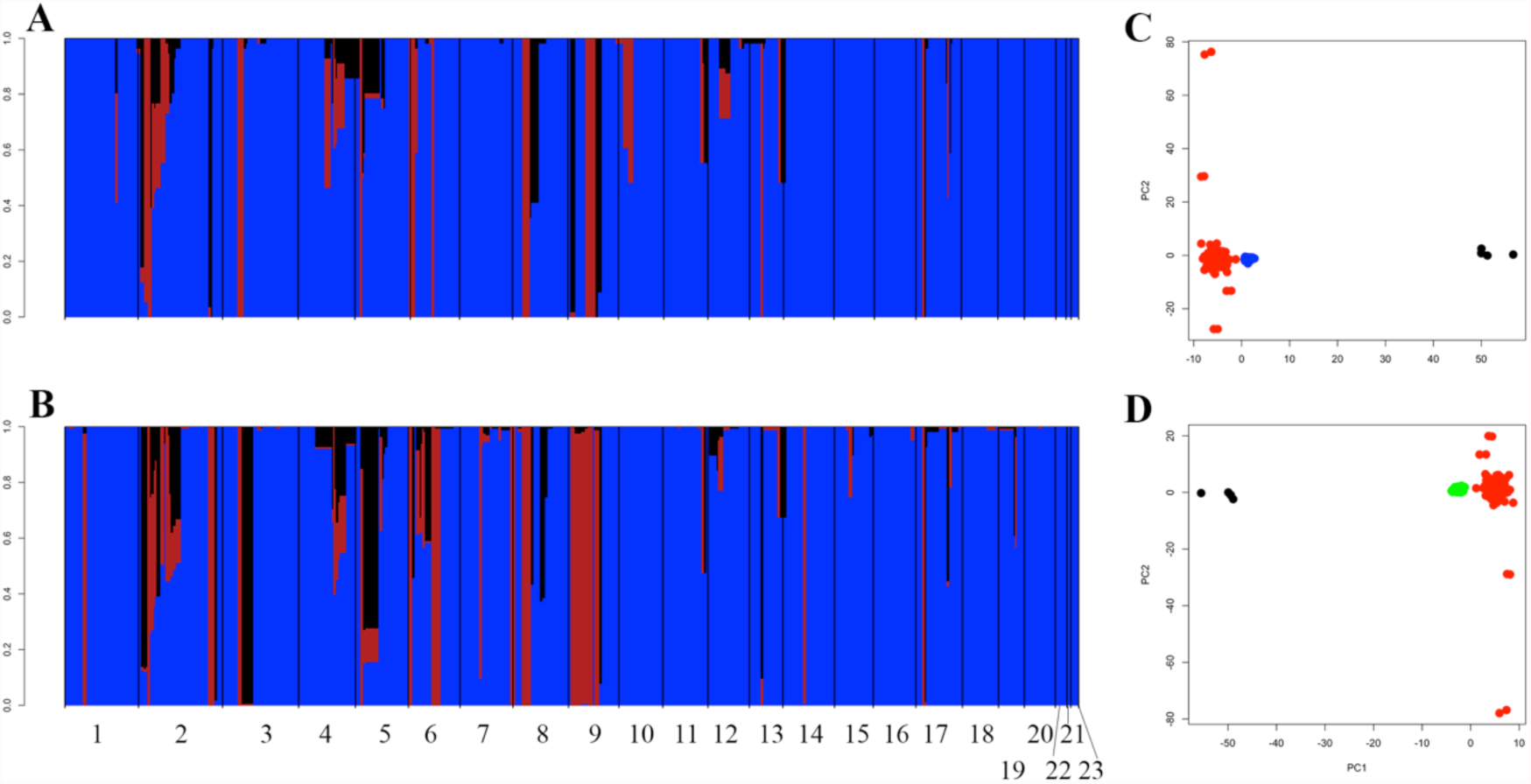
Local ancestry assignment of Estonian and Norwegian *H. fraxinues individuals*. The PCAdmix analyses give the average ancestry in windows of 10 SNPs along the genome (Russian ancestry in blue, Japanese ancestry in red and with uncertain ancestry in black) in (A)Estonian individuals and (B) Norwegian individuals. (C) and (D) show the average loadings for the first two principle components axes for (A) and (B) respectively (Russian haplotypes are in red, Japanese haplotypes are in black, Estonian haplotypes are in blue and Norwegian haplotypes are in green).

## Discussion

### Origin of European isolates of *H. fraxineus*

Genome-wide population diversity analysis using ddRAD allows increased sampling effort (increasing power), and provided important information on all stages of the invasion pathway including the introduction, the local establishment phase and the spread of *H. fraxineus* throughout Europe. The present study confirms the findings of McMullan et al., (submitted), that two parental individuals of *H. fraxineus* generated a self-sustaining population but in addition, the data presented here shows that these progenies then underwent a brief establishment phase and then swept throughout Europe in a short period of time. We suggest that *H. fraxineus* was introduced from the Russian Far East 14-32 years before the range expansion, which initiated probably around the time when the first symptoms occurred in the early 1990’s (Przybył 2002). Our results establish that *H. fraxineus* has a low introduction effort but also that no other effective secondary contact occurred during the range expansion process.

The Russian Far East ancestry of the European *H. fraxineus* was much more likely than an ancestry directly from Japan. In addition to Japan and the Russian Far East, *H. fraxineus* has been found on both *Fraxinus mandshurica* and *F. rhynchophylla* in South Korea (Han et al. 2014) and on *Fraxinus mandshurica* in China (Zheng and Zhuang 2014). Both the large number of fixed genomic regions, the lack of genetic structure and the presence of two haplotypes (McMullan et al (submitted)) show that *H. fraxineus* went through a severe genetic bottleneck upon introduction to Europe. Importantly we show that in the absence of secondary introductions since the founder event, 55% of the pathogen genome has become fixed and this signature is found throughout Europe. Effects of a founder event are such that alleles that are rare in the source population may be fixed upon introduction, through genetic drift. Such a scenario could explain the difference in the predominant ITS (internal transcribed spacer of rDNA) genotype between Europe and Asia (Drenkhan et al. 2017). Given the Japanese assignment of 10% - 15% of the European *H. fraxineus* genome, we cannot completely rule out that one isolate with a Japanese origin participated in the foundation of the European population. However, the fixed segments in the genome suggest that longer tracts with Japanese ancestry would probably be identified if this were the case. Thus it is much more likely that an introgression of Japanese ancestry occurred in the native population before introduction into Europe. Broader sampling of the native range of *F. mandshurica* and *F. rhynchophylla* may lead to the identification of a closer Asian source to European population, although this may be complicated by the signal of high gene flow in eastern Asia. Similar genetic structure was detected with microsatellites in Japan (Gross et al. 2014). Drenkhan et al. (2014) found evidence for import of the Russian *Fraxinus mandshurica* from the Far East into Baltic countries during Soviet occupation, supporting this as a possible route of introduction. In particular, *Fraxinus mandshurica* seeds and plants were imported to Estonia from the 1960s to 1980s. *H. fraxineus* has been identified on seeds on both *F. excelsior* and *F. mandshurica* using species specific primers (Cleary et al. 2013, Drenkhan et al. 2017), suggesting seeds as a possible path of introduction. Although the pathogen DNA is present in the seeds it has proved difficult to isolate the fungus from this material (Drenkhan, pers communication). Even if *H. fraxineus* may have been spread by seeds, the highest risks, and most obvious route, would be through the movement of plant material such as seedlings and wood that support sexual or asexual proliferation of the fungus (Husson et al. 2012, Gross et al. 2014). Based on the length of the diversity islands (or fixed blocks), we estimated the introduction of *H. fraxineus* occurred 14-32 years before the range expansion in Europe started, which overlaps well with the period of importation for *F. mandshurica* (Drenkhan et al. 2014). While this estimate is reliant on recombination rate, we believe that the recombination rate from *Botrytis cinerea* is probably closer to the expected rate in *H. fraxineus*, since the species both belong to the fungal order Helotiales. This suggests a scenario where as few as two haplotypes of *H. fraxineus* arrived in Europe in the 1970’s or the early 1980’s, when the Baltic States were still a part of the Soviet Union and the population increased locally before expansion outside the area of introduction.

Although there has been a dramatic change in genetic diversity during the founder event a relatively large amount of variation has been retained within the diverse regions. McMullan et al (submitted) found that part of the diversity of effector genes was preserved in the European population due to the introduction of two divergent haplotypes. Similarly, we found relatively high numbers of loci were outlier loci that are potentially under selection in the different regions in Europe, suggesting that *H. fraxineus* has further potential for local adaptation. Further studies are needed to ascertain the function of the regions under selection. Local adaptation to differences in climate were detected in recent range expansions of *Cryphonectria parasitica* in France; isolates from northern France showed growth optima at lower temperatures than those from southern France signifying that local adaptation evolved in a relatively short time (Robin et al. 2017).

The native European *Hymenoscyphus albidus,* which is a harmless leaf associate of European Ash (Baral and Bemmann 2014) could serve as a source of new genetic variation if hybridization occurs. However, no hybridization occurred in inter-specific crosses and *H. albidus* appears to reproduce exclusively via haploid selfing (Wey et al. 2016). These results are congruent with our genome-wide analysis which shows no signs of sexually or parasexually mediated introgression from *H. albidus* into the *H. fraxineus* genome. Our preliminary data on the genetic diversity of *H. albidus* suggests that the predominant mode of reproduction is selfing since very little variation was observed in the examined isolates from both Norway and Switzerland. More likely is that the niche-specialist *H. albidus* is just being solely outcompeted for the same sporulation niche and vastly overwhelmed by the mass of airborne *H. fraxineus* ascospores (Hietala et al. 2013).

### Biosecurity of *H. fraxineus*

The European pathogen population is bottlenecked and so movement of the host within Europe is unlikely to impact the allele frequencies. A real concern is a novel immigration event; even the introduction of a single isolate from outside of the European area, from locations such as Japan and Far Eastern Russia, could dramatically alter the genetic diversity of the Ash pathogen. Predictive modelling and an assessment of the likelihood of arrival are required to understand the impact of this introgression into the European area. We would like to emphasize the role of sexual reproduction and gene flow as a mechanism to introduce immigrant variation into the European *H. fraxineus* population. Such a scenario might dramatically increase pathogen fitness and impose further selection pressure and a steeper decline on the European host.

Declining European Ash populations and further fragmentation of the natural ranges of Ash, as well as the presence of resistance in host range margins (Tollefsrud et al. 2016), may inhibit the establishment of new hybrid *H. fraxineus* types. Nonetheless, the European *H. fraxineus* invasive range is extensive, and expanding. Presently, bioclimatic niche models for invasive alien species are fundamental tools used for pest risk assessment (Guisan and Thuiller 2005, Kriticos et al. 2013). We believe that defining the genomic diversity of the invasive species in both the invaded and native ranges is similarly essential. The genetic diversity of this pathogen may already impact its ability to adapt to host selective pressures.

However, the pathogen’s success in Europe, given the extreme nature of the primary founder event, reflects the fragility of Ash ecosystem. The present analysis was assisted by a fast sensitive surveillance and detection of genome-wide variation by scans gathered using ddRAD sequencing. This work, along with a refinement in the identification of the source population, will enable biosecurity managers to prioritize and make key decisions over resource management and assets, and to mitigate damage from Ash dieback, as well as for breeding an Ash recovery and future pest risk assessments.

## Author contributions

JHS, AVS, AH and HS designed the experiment. AH, HS, RD and KA collected the field material. AVS, JHS, AH, KA and RD performed DNA extractions and sequencing. JHS, AVS performed bioinformatic analysis. JHS, AVS and AH wrote the manuscript and with revisions from all authors.

## Acknowledgments

Financial contribution has been given by The Norwegian Genetic Resource Centre, The Research Council of Norway (grants 203822/E40 and 235947/E40), the Norwegian Financial Mechanism 2009–2014 under the project EMP162, the Institutional Research Funding IUT21-04, EMU project P170053MIMK, COST Action 1103 FRAXBACK and the Ministry of Agriculture and Food (131001).

We thank Dr Mark McMullan, Dr Matthew D. Clark, the NORNEX and OADB consortia for constructive comments and for sharing genomic data used in this study. We also thank Olaug Olsen and Gro Wollebæk (NIBIO), the laboratory of Prof Makoto Kakishima (Tsukuba, Japan), and the CBS-KNAW Fungal Biodiversity Centre (Utrecht, The Netherlands), for culturing and isolates, and Anne B. Nilsen and Jørn Petter Storholt (NIBIO) for GIS assistance.

## References

Amselem, J., C. A. Cuomo, J. A. L. van Kan, et al. “Genomic Analysis of the Necrotrophic Fungal Pathogens *Sclerotinia sclerotiorum* and *Botrytis cinerea*.” PLOS Genetics 7: e1002230(2011).

Baral, H.-O. & M. Bemmann. “*Hymenoscyphus fraxineus* vs. *Hymenoscyphus albidus* – A comparative light microscopic study on the causal agent of European ash dieback and related foliicolous, stroma-forming species.” Mycology 5: 228–290 (2014).

Baral, H.-O., V. Queloz & T. Hosoya. “*Hymenoscyphus fraxineus*, the correct scientific name for the fungus causing ash dieback in Europe.” IMA Fungus 5: 79–80(2014).

Bengtsson, S. B. K., R. Vasaitis, T. Kirisits, et al. “Population structure of *Hymenoscyphus pseudoalbidus* and its genetic relationship to *Hymenoscyphus albidus*.” Fungal Ecology 5: 147–153(2012).

Blackburn, T. M., P. Pysek, S. Bacher, et al. “A proposed unified framework for biological invasions.” Trends in Ecology & Evolution 26: 333–339(2011).

Brisbin, A., K. Bryc, J. Byrnes, et al. “PCAdmix: Principal Components-Based Assignment of Ancestry along Each Chromosome in Individuals with Admixed Ancestry from Two or More Populations.” Human biology 84: 343–364(2012).

Browning, S. R. & B. L. Browning. “Rapid and accurate haplotype phasing and missing-data inference for whole-genome association studies by use of localized haplotype clustering.” American Journal of Human Genetics 81: 1084–1097(2007).

Burokiene, D., S. Prospero, E. Jung, et al. “Genetic population structure of the invasive ash dieback pathogen *Hymenoscyphus fraxineus* in its expanding range.” Biological Invasions 17: 2743–2756(2015).

Cleary, M. R., N. Arhipova, T. Gaitnieks, et al. “Natural infection of *Fraxinus excelsior* seeds by *Chalara fraxinea*.” Forest Pathology 43: 83–85(2013).

Cross, H., J. H. Sønstebø, N. E. Nagy, et al. “Fungal diversity and seasonal succession in ash leaves infected by the invasive ascomycete *Hymenoscyphus fraxineus*.” New Phytologist 213: 1405–1417(2017).

Danecek, P., A. Auton, G. Abecasis, et al. “The variant call format and VCFtools.” Bioinformatics 27: 2156–2158(2011).

Dawson, W., D. Moser, M. van Kleunen, et al. “Global hotspots and correlates of alien species richness across taxonomic groups.” Nature Ecology and Evolution 1: 0186(2017).

Drenkhan, R., T. Riit, K. Adamson, et al. “The earliest samples of Hymenoscyphus albidus vs. H. fraxineus in Estonian mycological herbaria.” Mycological Progress 15: 835–844(2016).

Drenkhan, R., H. Sander & M. Hanso. “Introduction of Mandshurian ash (*Fraxinus mandshurica* Rupr.) to Estonia: Is it related to the current epidemic on European ash (*F. excelsior* L.)?” European Journal of Forest Research 133: 769–781(2014).

Drenkhan, R., H. Solheim, A. Bogacheva, et al. “*Hymenoscyphus fraxineus* is a leaf pathogen of local *Fraxinus* species in the Russian Far East.” Plant Pathology 66: 490–500(2017).

Earl, D. A. & B. M. vonHoldt. “STRUCTURE HARVESTER: a website and program for visualizing STRUCTURE output and implementing the Evanno method.” Conservation Genetics Resources 4: 359–361(2012).

Evanno, G., S. Regnaut & J. Goudet. “Detecting the number of clusters of individuals using the software STRUCTURE: a simulation study.” Molecular Ecology 14: 2611–2620(2005).

Falush, D., M. Stephens & J. K. Pritchard. “Inference of population structure using multilocus genotype data: Linked loci and correlated allele frequencies.” Genetics 164: 1567–1587(2003).

Fauvergue, X., E. Vercken, T. Malausa, et al. “The biology of small, introduced populations, with special reference to biological control.” Evolutionary Applications 5: 424–443(2012).

Foll, M. & O. Gaggiotti. “A Genome-Scan Method to Identify Selected Loci Appropriate for Both Dominant and Codominant Markers: A Bayesian Perspective.” Genetics 180: 977–993(2008).

Garrison, E. & G. Marth Haplotype-based variant detection from short-read sequencing. <u>preprint arXiv 1207.3907</u> (2012).

Gonthier, P. & M. Garbelotto. “Amplified fragment length polymorphism and sequence analyses reveal massive gene introgression from the European fungal pathogen *Heterobasidion annosum* into its introduced congener *H. irregulare*.” Molecular Ecology 20: 2756–2770(2011).

Gross, A., C. R. Grünig, V. Queloz, et al. “A molecular toolkit for population genetic investigations of the ash dieback pathogen *Hymenoscyphus pseudoalbidus*.” Forest Pathology 42: 252–264(2012).

Gross, A., O. Holdenrieder, M. Pautasso, et al. “*Hymenoscyphus pseudoalbidus*, the causal agent of European ash dieback.” Molecular Plant Pathology 15: 5–21(2014).

Gross, A., T. Hosoya & V. Queloz. “Population structure of the invasive forest pathogen *Hymenoscyphus pseudoalbidus*.” Molecular Ecology 23: 2943–2960(2014).

Gross, A., P. L. Zaffarano, A. Duo, et al. “Reproductive mode and life cycle of the ash dieback pathogen *Hymenoscyphus pseudoalbidus*.” Fungal Genetics and Biology 49: 977–986(2012).

Grunwald, N. J., B. A. McDonald & M. G. Milgroom. “Population Genomics of Fungal and Oomycete Pathogens.” Annual Review of Phytopathology 54: 323–346(2016).

Guisan, A. & W. Thuiller. “Predicting species distribution: offering more than simple habitat models.” Ecology Letters 8: 993–1009(2005).

Han, J.-G., B. Shrestha, T. Hosoya, et al. “First Report of the Ash Dieback Pathogen *Hymenoscyphus fraxineus* in Korea.” Mycobiology 42: 391–396(2014).

Hietala, A. M., V. Timmermann, I. Børja, et al. “The invasive ash dieback pathogen *Hymenoscyphus pseudoalbidus* exerts maximal infection pressure prior to the onset of host leaf senescence.” Fungal Ecology 6: 302–308(2013).

Huson, D. H. & D. Bryant. “Application of Phylogenetic Networks in Evolutionary Studies.” Molecular Biology and Evolution 23: 254–267(2006).

Husson, C., O. Caël, J. P. Grandjean, et al. “Occurrence of Hymenoscyphus pseudoalbidus on infected ash logs.” Plant Pathology 61: 889–895(2012).

Jakobsson, M. & N. A. Rosenberg. “CLUMPP: a cluster matching and permutation program for dealing with label switching and multimodality in analysis of population structure.” Bioinformatics 23: 1801–1806(2007).

Kowalski, T. & O. Holdenrieder. “Pathogenicity of *Chalara fraxinea*.” Forest Pathology 39: 1–7(2009).

Kraj, W., M. Zarek & T. Kowalski. “Genetic variability of *Chalara fraxinea*, dieback cause of European ash (*Fraxinus excelsior* L.).” Mycological Progress 11: 37–45(2012).

Kriticos, D. J., L. Morin, A. Leriche, et al. “Combining a Climatic Niche Model of an Invasive Fungus with Its Host Species Distributions to Identify Risks to Natural Assets: *Puccinia psidii* Sensu Lato in Australia.” PLoS ONE 8: e64479(2013).

Leggett, R. M., R. H. Ramirez-Gonzalez, B. J. Clavijo, et al. “Sequencing quality assessment tools to enable data-driven informatics for high throughput genomics.” Frontiers in Genetics 4: 288(2013).

Lischer, H. E. L. & L. Excoffier. “PGDSpider: an automated data conversion tool for connecting population genetics and genomics programs.” Bioinformatics 28: 298–299(2012).

Lockwood, J. L., P. Cassey & T. Blackburn. “The role of propagule pressure in explaining species invasions.” Trends in Ecology & Evolution 20: 223–228(2005).

McKinney, L. V., L. R. Nielsen, D. B. Collinge, et al. “The ash dieback crisis: genetic variation in resistance can prove a long-term solution.” Plant Pathology 63: 485–499(2014).

McKinney, L. V., I. M. Thomsen, E. D. Kjær, et al. “Rapid invasion by an aggressive pathogenic fungus (*Hymenoscyphus pseudoalbidus*) replaces a native decomposer (*Hymenoscyphus albidus*): a case of local cryptic extinction?” Fungal Ecology 5: 663–669(2012).

McMullan, M., M. Rafiqi, G. Kaithakottil, et al. “The ash dieback invasion of Europe was founded by two individuals from a native population with huge adaptive potential.” Nature Ecology and Evolution(submitted).

Narum, S. R. & J. E. Hess. “Comparison of FST outlier tests for SNP loci under selection.” Molecular Ecology Resources 11: 184–194(2011).

Paini, D. R., A. W. Sheppard, D. C. Cook, et al. “Global threat to agriculture from invasive species.” Proceedings of the National Academy of Sciences 113: 7575–7579(2016).

Pfeifer, B., U. Wittelsbürger, S. E. Ramos-Onsins, et al. “PopGenome: An Efficient Swiss Army Knife for Population Genomic Analyses in R.” Molecular Biology and Evolution 31: 1929–1936(2014).

Pritchard, J. K., M. Stephens & P. Donnelly. “Inference of population structure using multilocus genotype data.” Genetics 155: 945–959(2000).

Przybył, K. “Fungi associated with necrotic apical parts of *Fraxinus excelsior* shoots.” Forest Pathology 32: 387–394(2002).

Robin, C., A. Andanson, G. Saint-Jean, et al. “What was old is new again: thermal adaptation within clonal lineages during range expansion in a fungal pathogen.” Molecular Ecology 26: 1952–1963(2017).

Roper, M., A. Seminara, M. M. Bandi, et al. “Dispersal of fungal spores on a cooperatively generated wind.” Proceedings of the National Academy of Sciences 107: 17474–17479(2010).

Rytkönen, A., A. Lilja, R. Drenkhan, et al. “First record of *Chalara fraxinea* in Finland and genetic variation among isolates sampled from Åland, mainland Finland, Estonia and Latvia.” Forest Pathology 41: 169–174(2011).

Saunders, D., K. Yoshida, C. Sambles, et al. “Crowdsourced analysis of ash and ash dieback through the Open Ash Dieback project: A year 1 report on datasets and analyses contributed by a self-organising community.” bioRxiv, 004564 (2014).

Schoebel, C. N., L. Botella, V. Lygis, et al. “Population genetic analysis of a parasitic mycovirus to infer the invasion history of its fungal host.” Molecular Ecology 26: 2482–2497(2017).

Skovsgaard, J. P., I. M. Thomsen, I. M. Skovgaard, et al. “Associations among symptoms of dieback in even-aged stands of ash (Fraxinus excelsior L.).” Forest Pathology 40: 7–18(2010).

Solheim, H. & A. M. Hietala. “Spread of ash dieback in Norway.” Baltic Forestry 23: 144–149(2017).

Stukenbrock, E. H., T. Bataillon, J. Y. Dutheil, et al. “The making of a new pathogen: Insights from comparative population genomics of the domesticated wheat pathogen *Mycosphaerella graminicola* and its wild sister species.” Genome Research 21: 2157–2166(2011).

Stukenbrock, E. H., F. B. Christiansen, T. T. Hansen, et al. “Fusion of two divergent fungal individuals led to the recent emergence of a unique widespread pathogen species.” Proceedings of the National Academy of Sciences 109: 10954–10959(2012).

Timmermann, V., I. Børja, A. M. Hietala, et al. “Ash dieback: pathogen spread and diurnal patterns of ascospore dispersal, with special emphasis on Norway.” EPPO Bulletin 41: 14–20(2011).

Tollefsrud, M. M., T. Myking, J. H. Sønstebø, et al. “Genetic Structure in the Northern Range Margins of Common Ash, *Fraxinus excelsior* L.” PLOS ONE 11: e0167104(2016).

Van Kan, J. A. L., J. H. M. Stassen, A. Mosbach, et al. “A gapless genome sequence of the fungus *Botrytis cinerea*.” Molecular Plant Pathology 18: 75–89(2017).

Wey, T., M. Schlegel, S. Stroheker, et al. “MAT--gene structure and mating behavior of *Hymenoscyphus fraxineus* and *Hymenoscyphus albidus*.” Fungal Genetics and Biology 87: 54–63(2016).

Zhan, A., J. A. Darling, D. G. Bock, et al. “Complex genetic patterns in closely related colonizing invasive species.” Ecology and Evolution 2: 1331–1346(2012).

Zhao, Y.-J., T. Hosoya, H.-O. Baral, et al. “*Hymenoscyphus pseudoalbidus*, the correct name for *Lambertella albida* reported from Japan.” Mycotaxon 122: 25–41(2013).

Zheng, H.-D. & W.-Y. Zhuang. “*Hymenoscyphus albidoides* sp. nov. and *H. pseudoalbidus* from China.” Mycological Progress 13: 625–638(2014).

Zheng, X., D. Levine, J. Shen, et al. “A high-performance computing toolset for relatedness and principal component analysis of SNP data.” Bioinformatics 28: 3326–3328(2012).

